# Cross-protein transfer learning substantially improves disease variant prediction

**DOI:** 10.1101/2022.11.15.516532

**Authors:** Milind Jagota, Chengzhong Ye, Carlos Albors, Ruchir Rastogi, Antoine Koehl, Nilah Ioannidis, Yun S. Song

**Affiliations:** Computer Science Division, University of California, Berkeley, CA 94720; Department of Statistics, University of California, Berkeley, CA 94720; Chan Zuckerberg Biohub, San Francisco, CA 94158; Center for Computational Biology, University of California, Berkeley, CA 94720

## Abstract

Genetic variation in the human genome is a major determinant of individual disease risk, but the vast majority of missense variants have unknown etiological effects. Here, we present a robust learning framework for leveraging saturation mutagenesis experiments to construct accurate computational predictors of proteome-wide missense variant pathogenicity. We train cross-protein transfer (CPT) models using deep mutational scanning data from only five proteins and achieve state-of-the-art performance on clinical variant interpretation for unseen proteins across the human proteome. High sensitivity is crucial for clinical applications and our model CPT-1 particularly excels in this regime. For instance, at 95% sensitivity of detecting human disease variants annotated in ClinVar, CPT-1 improves specificity to 68%, from 27% for ESM-1v and 55% for EVE. Furthermore, for genes not used to train REVEL, a supervised method widely used by clinicians, we show that CPT-1 compares favorably with REVEL. Our framework combines predictive features derived from general protein sequence models, vertebrate sequence alignments, and AlphaFold2 structures, and it is adaptable to the future inclusion of other sources of information. We find that vertebrate alignments, albeit rather shallow with only 100 genomes, provide a strong signal for variant pathogenicity prediction that is complementary to recent deep learning-based models trained on massive amounts of protein sequence data. We release predictions for all possible missense variants in 90% of human genes. Our results demonstrate the utility of mutational scanning data for learning properties of variants that transfer to unseen proteins.

## Introduction

Variation in the human genome across individuals is a major determinant of differences in disease risk, and exponential decreases in sequencing costs have made it feasible to measure personal genome sequences of individual patients. To be able to make accurate and targeted medical decisions based on genetic information, we need to understand the etiological consequences of human genome variants. Missense variants, which modify the amino acid at a single position of a protein, are of particular interest because their effects on protein structure and function are highly variable. Tens of millions of missense variants may exist in the human population, and the vast majority of these have unknown consequences [1, 2]. Efforts to collect population genomic data relating variants to disease phenotypes have made progress on this problem [3, 4]. However, many of the missense variants in the human population only exist in a tiny fraction of individuals and may not clearly present their disease consequences, thus limiting the ability of population genomic data to inform variant effects. Functional assays via deep mutational scanning (DMS) experiments have been used to measure the effects of missense variants at higher throughput [5–7]. However, these approaches still do not directly scale to the whole human proteome, and depend on the ability to design a relevant assay for each protein of interest.

There has also been significant interest in developing computational methods to predict the effects of missense variants [8–15]. Computational methods can provide predictions for all possible mutations across the human proteome and have proven to be effective predictors of variant pathogenicity. Recently, the methods EVE [8] and ESM-1v [9] have been demonstrated to achieve state-of-the-art performance in human disease variant prediction and functional assay prediction, respectively. EVE and ESM-1v achieve strong performance despite not training on human clinical data or functional assays. Instead, the underlying principle of these models is to collect large databases of natural protein sequences, then model the probability distribution of how protein sequences vary. However, these models also have limitations. EVE and ESM-1v model protein sequence variation across all known species, and employ redundancy filtering so that variation between highly similar protein sequences (such as from related species) is not considered. This approach allows these methods to effectively capture broad constraints of protein families, such as those imposed by a common structural fold [16–20]. While this signal is powerful, it is not sufficient to fully explain how an amino acid substitution impacts protein function. This limitation has been shown by several recent studies in the context of functional assay prediction, which have found that the accuracy of sequence variation methods is far from the reproducibility of functional assays [21–23]. Moreover, sequence variation methods can be significantly improved by learning on functional data for a specific system of interest [21, 22].

Analagous to these results in functional assay prediction, we postulated that protein sequence models such as EVE and ESM-1v could be combined with more human-specific sources of information to improve their performance on variant pathogenicity prediction. The most widely-used ensemble models have used training data derived from clinical variant annotations or population genomic information [10–13]. However, such models can be affected by circularity and bias in the collection of these data, such as the use of other computational predictors to generate variant annotations [24, 25]. To produce a broadly applicable model that would generalize to diverse target proteins, we instead developed a robust learning framework to train disease variant predictors using functional assay data from a small number of proteins (Figure 1). Functional assay data measure many variants per protein, allowing us to obtain sufficient data from very few proteins while saving the vast majority of the human proteome as a fully unseen evaluation. These data are also typically exhaustive (as in DMS) or generated with a well-defined process for choosing variants, avoiding biases of clinical data. In our work, we trained cross-protein transfer (CPT) models using DMS data from only five proteins, all from the same functional assay relevant to human pathogenicity, and achieved significantly improved performance over EVE and ESM-1v on clinical variant interpretation. Furthermore, for genes not used to train REVEL, an ensemble method widely used by clinicians, we demonstrate that our model CPT-1 compares favorably with REVEL. We therefore expect that our predictions are accurate and more robust than what has previously been available, and we publicly release predictions for all possible missense variants in 90% of human genes. Previous work has trained on DMS data [26, 27], but these models did not match the performance of those supervised on clinical data.

**Figure 1:**
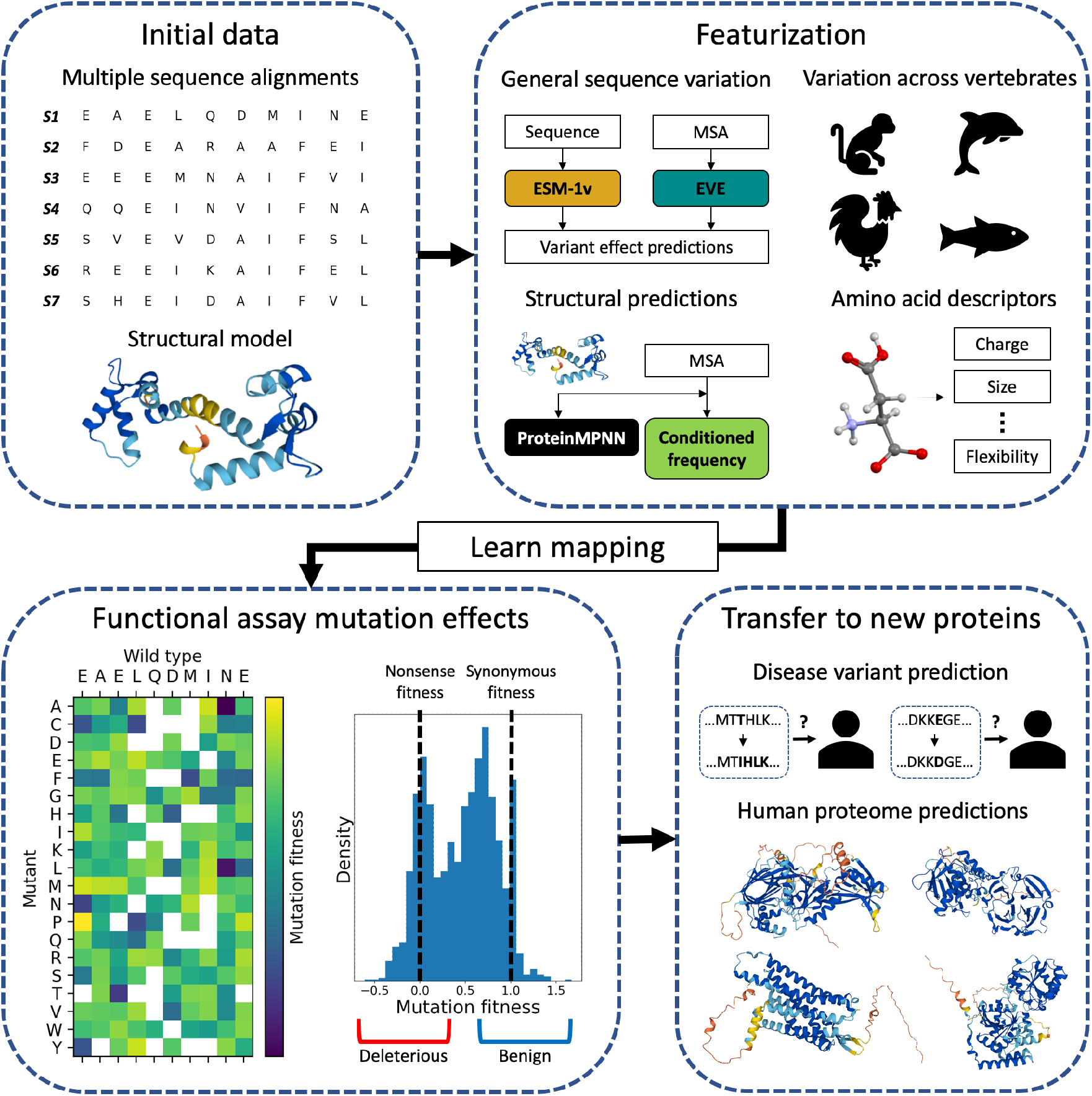
Method Overview. We develop computational missense variant effect predictors by training on functional assay data from very few proteins and achieve substantially improved performance over the state-of-the-art. We combine general protein sequence variation (EVE, ESM1v), sequence variation at local evolutionary timescales (vertebrate alignments), protein structure (AlphaFold2, ProteinMPNN), and amino acid representations. We assess our models on unseen proteins across the human proteome and release predictions for all missense variants in 90% of human genes.

Our model integrates features from multiple sequence alignments (MSAs) at local evolutionary timescales and explicit protein structure models together with state-of-the-art zero-shot variant effect predictors such as EVE and ESM-1v. To prevent data leakage, we did not use any features which were previously trained on clinical or functional assay data of other proteins. For our MSA features, we used alignments of 100 vertebrates and 30 mammals which were constructed using whole-genome alignment, providing small collections of orthologous sequences that have a higher degree of functional conservation compared to the data used by EVE and ESM-1v [28, 29]. Also, we leveraged AlphaFold2 structure models to provide the specific structure of proteins in a representation that allows use of features based on protein geometry [30, 31]. Our framework is adaptable to the future inclusion of other predictors. While functional assay data do not readily scale to the whole proteome directly, our results demonstrate the utility of relatively small amounts of such data for enhancing computational predictors of disease variants.

## Results

### State-of-the-art accuracy on clinical variants and functional assays

We trained a model, CPT-1, to classify missense variants as benign or pathogenic, using only DMS data from five human proteins (Figure 1, Methods). These proteins (CALM1, MTHR, SUMO1, UBC9, and TPK1) were studied using the same fitness assay by the same lab [7,32], which provided a controlled, high-quality training dataset. We also experimented with training on additional human DMS datasets and discuss the results of these experiments later in this section. CPT-1 integrates the general protein sequence models EVE [8] and ESM-1v [9] with conservation features from vertebrate alignments and structural features calculated using AlphaFold2 structures. Starting from a large list of candidate features, we performed feature selection using cross validation on DMS data and selected nine features for the final model (Supplementary Table S1, Methods).

We assessed CPT-1 for clinical disease variant prediction using ClinVar missense variants in human genes that are annotated as benign or pathogenic [1] (Figure 2A-C, Supplementary Table S2). We used all ClinVar variants released from 2017 onward that have at least a one star annotation, and also restricted to genes where EVE scores are available [8]. This left us with a high-quality dataset of 24,155 variants in 1298 genes (Methods). We primarily report comparisons to EVE and ESM-1v which, like CPT-1, do not train on clinical or functional assay data from evaluation proteins. EVE has been comprehensively evaluated against other well-known methods and shown to achieve competitive or superior performance [8]. We additionally compare our model with REVEL, an ensemble method widely used by clinicians [10]. REVEL is supervised on clinical variant annotations. Hence, to ensure a fair comparison, we constructed a separate dataset from which we remove all genes that had a clinical variant annotation available at the time that REVEL was trained (Methods). This dataset is rather small (3754 variants in 407 genes) and we therefore focus primarily on our full ClinVar test set.

**Figure 2:**
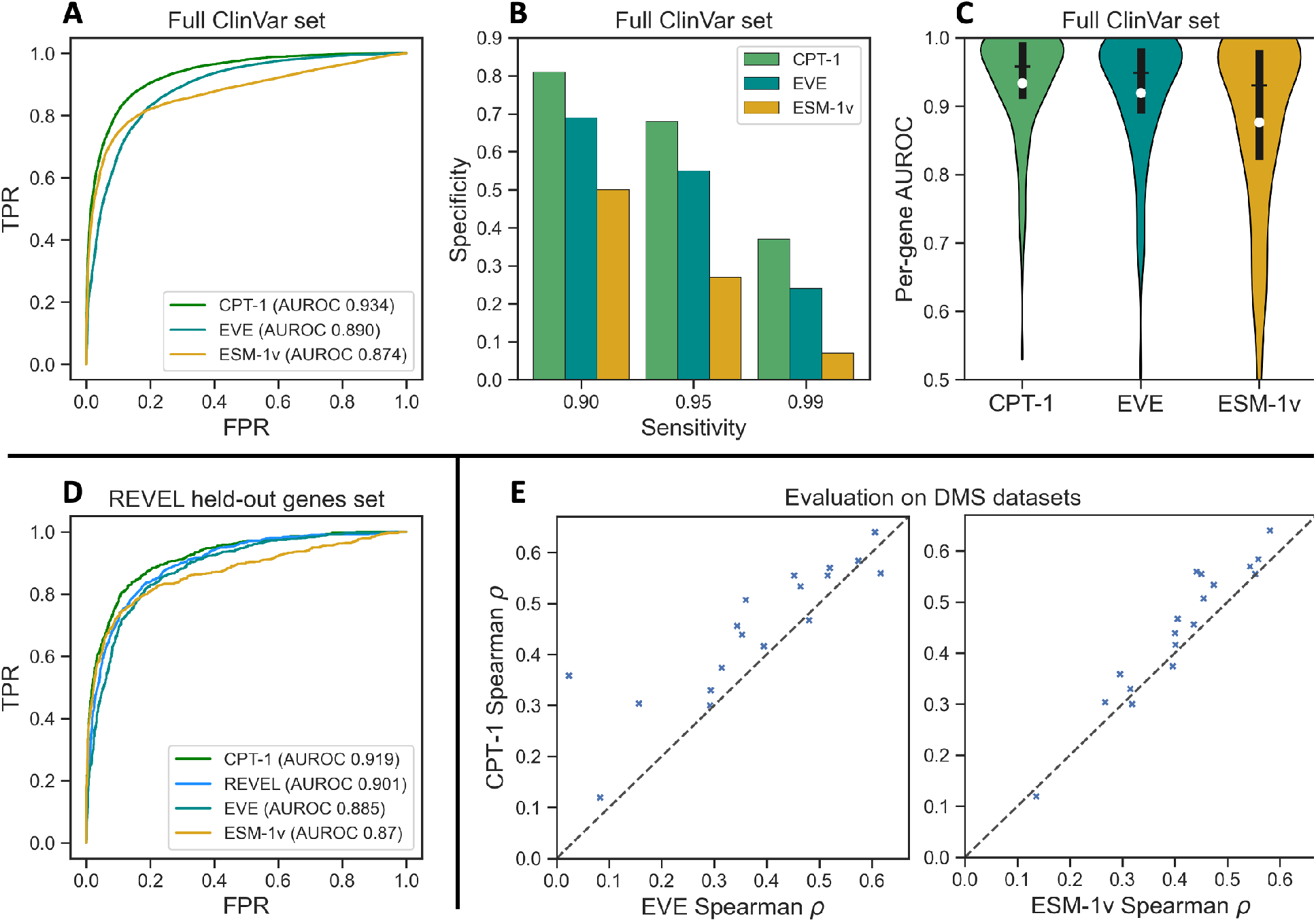
CPT-1 achieves state-of-the-art performance on clinical variant and functional assay prediction. (A) Receiver-operating characteristic (ROC) curves for ESM-1v, EVE, and our transfer model CPT-1 on annotated missense variants in ClinVar. CPT-1 improves the true positive rate at all false positive rates over both baselines and has a significantly higher AUROC. (B) Specificity in the clinically relevant high-sensitivity regime on ClinVar missense variants. When all models are constrained to recall nearly all pathogenic variants, CPT-1 improves on EVE and ESM-1v by large margins. (C) Per-gene AUROC on ClinVar missense variants in 886 genes with at least four benign and four pathogenic variants. Interquartile range and median shown in black, mean shown in white. CPT-1 improves or equals the per-gene AUROC on 72% of genes for EVE and 79% of genes for ESM-1v. (D) CPT-1 outperforms REVEL on proteins that were not trained on by REVEL, demonstrating the value of developing predictors with cross-protein transfer in mind. (E) Spearman’s *ρ* on DMS datasets of human proteins from ProteinGym (full details in Supplementary Table S3). The left plot compares CPT-1 to EVE, and the right compares CPT-1 to ESM-1v. In each plot, points above the diagonal line indicate a gene where CPT-1 outperforms the baseline. With the test protein held out in all cases, CPT-1 outperforms EVE on 16 out of 18 proteins and outperforms ESM-1v on 15 out of 18.

CPT-1 achieves substantially improved performance over EVE and ESM-1v, despite not training directly on any proteins in our assessment dataset. CPT-1 has an improved sensitivity (or true positive rate) for any given specificity (or 1 – false positive rate) over both EVE and ESM-1v, and significantly improves the overall area under ROC curve (AUROC) (Figure 2A). Performance increases are particularly large in the clinically relevant high-sensitivity regime (Figure 2B), where a good computational predictor is expected to flag almost all pathogenic variants with as high specificity (or as few false positives) as possible [15]; for example, at 95% sensitivity, CPT-1 improves specificity to 68%, from 27% for ESM-1v and 55% for EVE. Out of 13,815 benign variants in our dataset, this corresponds to nearly 1,800 fewer false positives compared to EVE and over 5,500 fewer false positives compared to ESM-1v when classifying 95% of pathogenic variants correctly. We also examined per-protein performance in our dataset for those proteins with at least four benign and four pathogenic Clinvar missense variants (Figure 2C, Supplementary Table S2). CPT-1 achieved improved or equal AUROC compared to EVE and ESM-1v on 72% and 84% of genes, respectively (strictly greater AUROC on 53% and 61% of genes, respectively). In our REVEL held-out genes set, CPT-1 outperforms REVEL as well as ESM-1v and EVE (Figure 2D). If we additionally restrict to rare variants, the margin of CPT-1 over REVEL increases (Supplementary Figure S1). Compared to REVEL, CPT-1 has the additional utility of providing predictions for all possible amino acid variants and not just observed single nucleotide variants, and also relies on significantly fewer features.

We note that across all assessments, EVE has a higher per-gene AUROC than global AUROC. This is likely because EVE fits a separate density model for each gene of interest. This means that predictions are well-calibrated within each gene but it is difficult to pick up differences in average variant effect between genes. CPT-1 improves the per-gene AUROC of EVE and does not show the same relative gap to global AUROC, indicating that it has also captured differences in the average variant effect between genes.

We also assessed our cross-protein transfer framework for zero-shot functional assay (DMS) prediction (Figure 2E, Supplementary Table S3). We combined our five training datasets with 13 additional DMS datasets of human proteins from ProteinGym [33]. We then generated variant effect predictions for each protein, using a regression model that was trained only on other proteins (Methods). Our method achieves a higher Spearman’s *ρ* than EVE in 16 out of 18 proteins, and outperforms ESM-1v on the same metric in 15 out of 18 proteins. In total, CPT-1 is the outright best performer in 13 out of 18 proteins.

To conclusively establish our claims about the value of supervising on DMS, we compared the full CPT-1 model with several additional baselines (Figure 3). First, we compared the performance of CPT-1 with alternatives that do not rely on the DMS data as much (Figure 3A). Specifically, we compared CPT-1 with unweighted averaging of EVE and ESM-1v, unweighted averaging of randomly selected features, and unweighted averaging of the features selected by our feature selection procedure. CPT-1 outperforms all of these alternatives, especially in the clinically relevant highsensitivity regime. In particular, unweighted averaging of DMS-selected features performs worse than averaging ESM-1v and EVE, indicating that training on DMS goes beyond selecting features; the learned coefficients are essential to high performance.

**Figure 3:**
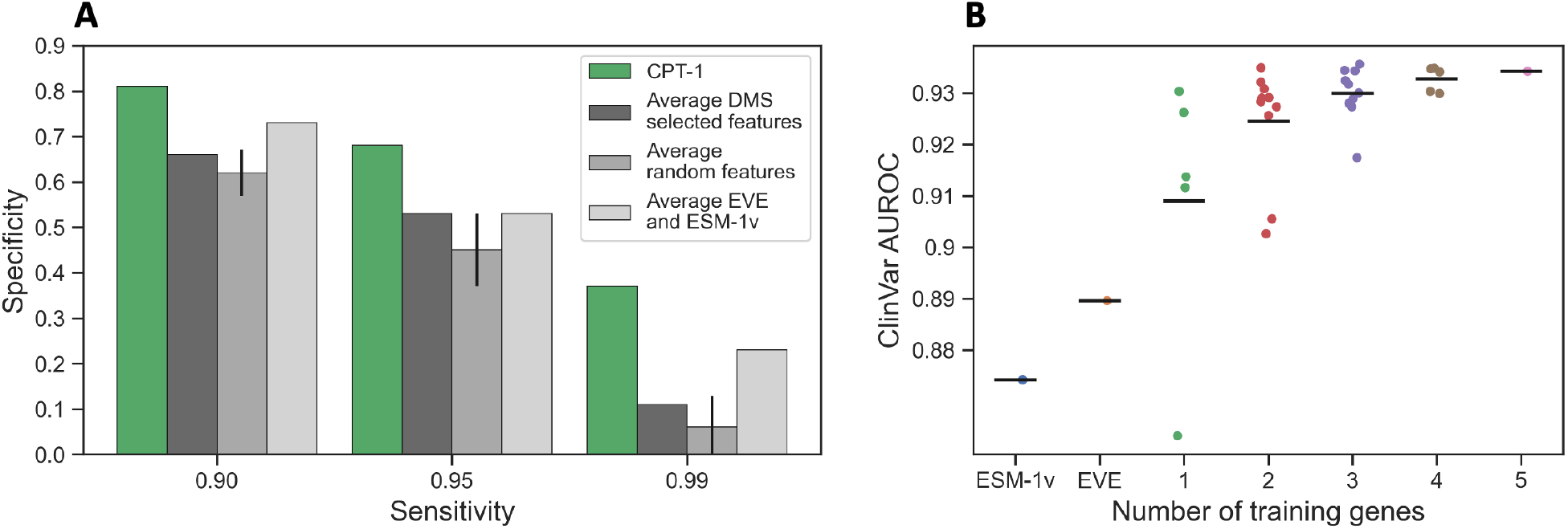
Training on DMS is important for CPT-1 performance. (A) We compared CPT-1 performance to several baselines that do not fully use the DMS data. These baselines were: averaging EVE and ESM-1v, averaging random features (set to the correct sign), and averaging features selected by feature selection. CPT-1 outperforms these baselines, especially in the high-sensitivity regime. This demonstrates the value of a full training procedure on DMS data. (B) We examined the dependence of CPT-1 performance on the number of training genes used. Each dot indicates a specific choice of training genes, with the mean shown as a black horizontal bar. More training genes always increases average performance, but there is significant variance and performance increases appear to be saturating. We also examined the use of additional, more heterogeneous datasets from ProteinGym, finding that this did not increase performance (Supplementary Figure S2).

We also measured the impact of the number of training genes used to train CPT-1 (Figure 3B). We found that average performance increases with the number of training genes and appears to be saturating at all five used. We additionally tested training on the aforementioned additional 13 human protein datasets in ProteinGym (Supplementary Figure S2). We found that our five chosen proteins from the same lab generally yielded higher performance than five random human proteins from ProteinGym, demonstrating the utility of using more consistent data. Moreover, training on all the human DMS datasets in ProteinGym did not improve performance beyond the five high quality datasets.

We note that EVE declines to make predictions on variants at positions where alignment quality is low, which make up 15% of variants in the ClinVar assessment dataset. We imputed EVE predictions at these variants using a nearest-neighbors approach within each gene (Methods). EVE scores are less accurate at these imputed positions but are still high-performing and improve model performance (Supplementary Figure S3). Also, Frazer *et al*. [8] reported performance for EVE with low confidence predictions removed. In our assessments, we include all predictions for all models. ESM-1v, meanwhile, does not accept proteins longer than 1,022 amino acids by default. We developed and implemented a scheme to apply ESM-1v to longer proteins, which make up 44% of the genes in our evaluation dataset (Methods). ESM-1v scores perform worse on these long genes, but EVE and CPT-1 do not suffer a loss in performance (Supplementary Figure S4).

### Vertebrate alignments are key to improved performance

EVE and ESM-1v achieve impressive performance using only protein sequence variability at the scale of the whole tree of life. Concretely, these models collect protein sequences from the set of all known proteins and employ redundancy filtering for sequences with high similarity. This approach models broadly recurring constraints well, such as structural constraints of a fold. However, we postulated that the models may be disregarding useful signal about sequence variation in species that are closer to humans.

We integrated two sets of alignments into CPT-1 to address this gap, extracted from 100 vertebrates and 30 mammals via whole-genome alignment, referred to generally as vertebrate multiple sequence alignments (vtMSA) (Methods) [28,29,34]. These alignments are shallow, but provide sequences that are orthologous to the target human protein and from species that are close to humans in the context of the full tree of life. The protein sequences are also often within the redundancy filtering criteria of EVE and ESM-1v. For example, EVE downweights sequences that are within 80% sequence identity of each other, but the average 100 vertebrate MSA has 44 sequences that are within 80% sequence identity of the human protein. Our features treat each of these 44 sequences as a full observation, whereas EVE downweights them to have a total weight of one observation. Likewise, ESM-1v uses sequences clustered at 90% identity, but the average 100 vertebrate MSA has 28 sequences that are within 90% identity of the human protein. These traits mean that conservation in these alignments is likely to be non-redundant with EVE and ESM-1v while being more specific to function and organism. Conservation in vertebrate alignments has previously been studied as a predictor of variant effects [35–37]; one of our main contributions is to show that this signal is useful even in the presence of much more powerful sequence variation methods.

Simple features from vertebrate alignments are competitive with models like EVE and ESM-1v for predicting of the pathogenicity of human disease variants (Figure 4A-B). The frequency of the wild-type amino acid in the aligned column of the 100-vertebrate MSAs, for example, achieves a global ClinVar AUROC of 0.865. This is close to the performance of ESM-1v and EVE, and better than single conservation features calculated from the much larger EVE MSA. In particular, 100-vertebrate wild-type frequency alone is competitive with EVE and ESM-1v in the high-sensitivity regime (Figure 4A). However, 100-vertebrate wild-type frequency has a lower overall AUROC than EVE and ESM-1v, and is much worse in the high specificity regime (Supplementary Table S2).

**Figure 4:**
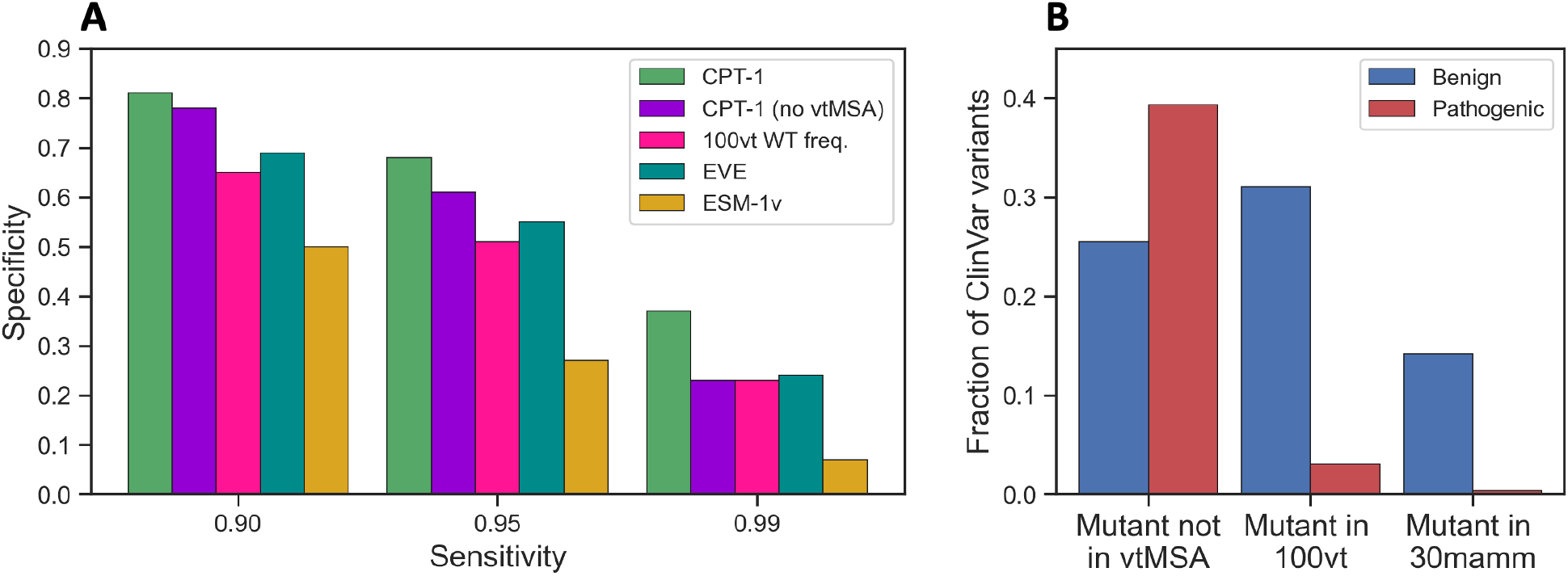
Vertebrate alignments are key to improved performance and a powerful baseline. (A) Specificity in the clinically relevant high-sensitivity regime on ClinVar missense variants. Removing vertebrate alignments from CPT-1 significantly decreases the margin of improvement over baseline. Conservation among 100 vertebrates is a powerful single feature baseline, and is competitive with much more complex models in the high-sensitivity regime. Vertebrate alignments are much less powerful in the high specificity regime (Supplementary Table S2). (B) If a missense variant from ClinVar appears in a vertebrate alignment, it is highly likely to be benign. Of the variants that do not occur in any of our studied vertebrates, 22% are benign. Of the variants that occur in a vertebrate, 82% are benign. Of the variants that occur in a mammal (subset of vertebrates), 94% are benign. This signal is not fully leveraged by EVE and ESM-1v due to the sequence redundancy filtering that is employed by both methods, and is key to the improved performance of CPT-1.

Furthermore, if a variant occurs at all in either the 100-vertebrate or 30-mammal MSA, it can be inferred to be benign with high probability (Figure 4B). Concretely, mutant (more precisely, non-reference in human) alleles that appear at the same position in the reference genome for at least one other vertebrate are 82% benign, and those that appear for at least one other mammal are 94% benign. In contrast, variants that are not the reference allele in any other vertebrate are 61% pathogenic. This aligns with the clinical practice of expecting human mutations with high allele frequency to be benign. A benign mutant allele can be at low frequency in humans, but its presence in 30-mammal or 100-vertebrate alignments suggests high frequencies in the corresponding species carrying the allele. This provides support for non-pathogenicity since these species are similar to humans in the context of the entire tree of life.

We also measured the importance of vertebrate alignments by training a CPT model that does not use them (Methods). This model performs substantially worse than our full model on clinical data, especially at the highest sensitivities (Figure 4A, Supplementary Table S2). This margin can be partially explained in terms of the feature presented in Figure 4B regarding frequency of variants in the vertebrate alignments. Suppose we set both EVE and CPT-1 to predict variants with a sensitivity of 99%. If a variant is predicted as pathogenic by EVE but appears as the reference allele for a non-human vertebrate, CPT-1 predicts it as pathogenic only 54% of the time. In contrast, if a variant is predicted as pathogenic by EVE and does not appear as the reference allele for a non-human vertebrate, CPT-1 predicts it as pathogenic 99% of the time. CPT-1 could make an incorrect prediction for variants that are pathogenic in humans but appear as the reference allele in another vertebrate, but we find that very few such variants exist.

At a per-gene level, adding vertebrate alignments is neutral or beneficial to the performance of CPT-1 in 84% of genes. The genes where vertebrate alignments help the most (top 10%) are more challenging genes in general, with lower average AUROCs for CPT-1, ESM-1v, and EVE. In addition, they have more shallow MSAs (average depth of 6,600 compared to 10,000 in all genes) and are longer (68% are longer than 1000 amino acids, compared to 44% in all genes). These discrepancies suggest that vertebrate alignments may be more useful in more complex human genes, which are more recently evolved and harder to model through general sequence homology.

### Insights from AlphaFold structures

Structural features provide a direct representation of protein geometry that can be informative of function. We used AlphaFold2 predicted structures from the AlphaFold2 human proteome database for all proteins in this study (Methods) [31]. There has been considerable interest in using AlphaFold2 structures for missense variant effect prediction [38–41]. We tested two major classes of features. First, we included multiple versions of the deep neural network ProteinMPNN (which takes structure as input) [42]. Second, we included two hand-designed features that combine a known structure with conservation in the EVE MSA. For the latter features, we aimed to compute wild-type and mutant frequencies conditioned on the structural environment being the same as in the human protein. To achieve this, we first find for each position all other positions which are in contact in the AlphaFold structure. We then filter the EVE MSA to sequences where these positions have the same amino acids as in the human sequence. However, we perform this filtering using only a maximum of two contact residues, to ensure the number of sequences does not become too small. We define these features precisely in the Methods.

Structural features slightly improve performance of CPT-1 (Figure 5A, Supplementary Table S2, Methods). These performance increases hold even though ProteinMPNN, which depends the least on sequence variability out of our major features, has low accuracy on its own. AlphaFold2 structures thus appear to encode useful information that is not captured from sequence variation alone. However, improvements from adding structure are much smaller than from adding vertebrate alignments. This is consistent with previous results showing that large protein sequence variability methods like EVE and ESM-1v model protein structure implicitly [16–19]. AlphaFold2 structures can be retrieved and analyzed very rapidly, which is an advantage over the extremely slow process of MSA generation. However, AlphaFold2 uses MSAs itself to make predictions; the release of these MSAs at scale would likely enable further increases in performance.

**Figure 5:**
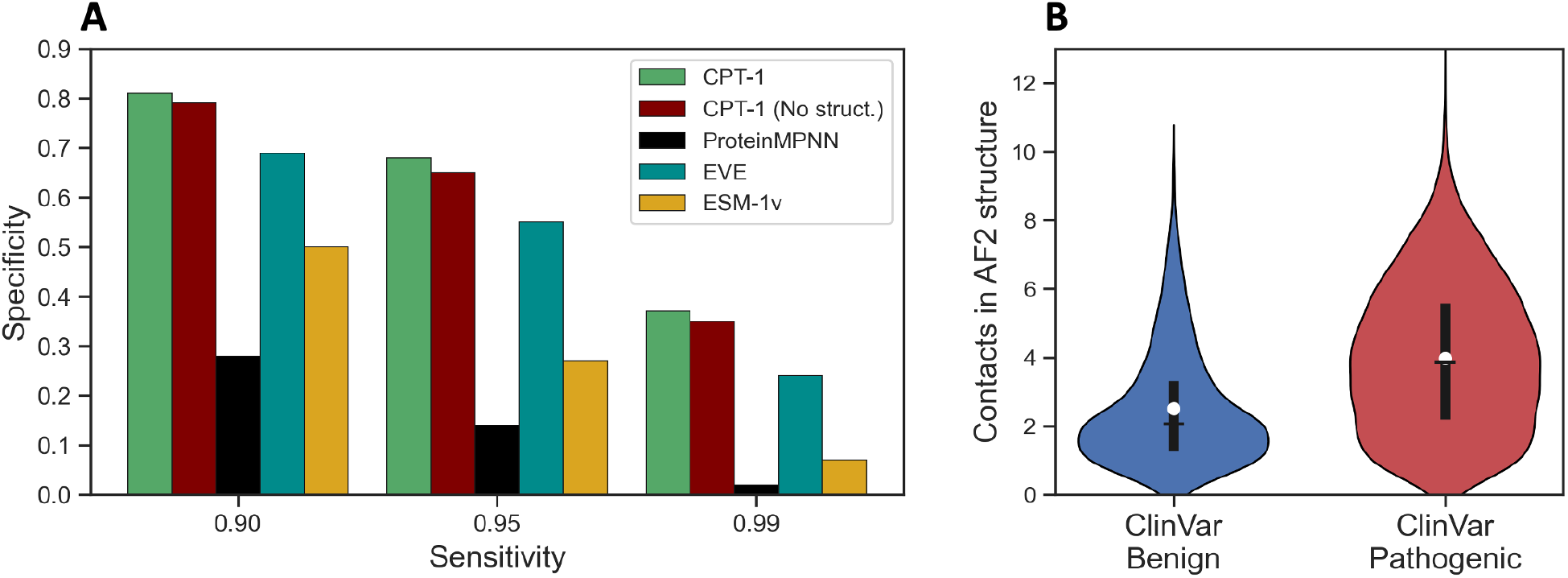
Insights from AlphaFold structures. (A) Specificity of CPT-1 in the clinically relevant high-sensitivity regime on ClinVar missense variants. Structural features slightly improve CPT-1 performance even though ProteinMPNN alone has poor performance. (B) Pathogenic ClinVar variants are more likely to have many contacts in the AlphaFold2 structure for the protein compared to benign variants.

The AlphaFold2 structural models for our five training proteins are all high-quality and informed by experimental structures, while many proteins in our clinical dataset do not have high quality AlphaFold2 models or have disordered regions. We analyzed the correlation of AlphaFold2 structure quality, as measured by AlphaFold pLDDT, with the performance of structural features in disease variant prediction but did not observe any significant signal. Even the performance of ProteinMPNN, which is trained exclusively on experimental protein structures, does not deteriorate dramatically when it is applied to structures with disordered or poorly modeled regions.

We observed that structural features of the site of variant such as contact count and AlphaFold pLDDT are directly predictive of variant pathogenicity (AUROC 0.69 for both), indicating that ClinVar variants in the structural cores of proteins are more likely to be pathogenic (Figure 5B). However, we found these features to be redundant with ProteinMPNN, indicating that Protein-MPNN captures this signal already. In addition, ProteinMPNN performs better on genes where ClinVar variant positions have more contacts in the AlphaFold2 structure (Methods, Supplementary Figure S5).

### Predictions across the human proteome

We looked to produce predictions from our method at whole-proteome scale. EVE MSAs and predictions are not available for the vast majority of human genes and are highly computationally intensive to compute. We therefore imputed all features that depend on the EVE MSA across genes using a nearest-neighbors approach (Supplementary Table S1, Methods). Then, using the aforementioned five functional DMS datasets, we refit coefficients of CPT-1 for use on cross-gene imputed features. We assessed this version of CPT-1 on our full ClinVar dataset and found that the model still outperformed EVE and ESM-1v (Figure 6A-B). We additionally compared CPT-1 to CPT-1 with imputed EVE and CPT-1 with no EVE (Figure 6C-D). Imputation improves performance compared to removing EVE entirely, but there is still a gap to having true EVE scores computed. This indicates that it will be useful to generate high quality MSAs and EVE predictions for the entire human proteome.

**Figure 6:**
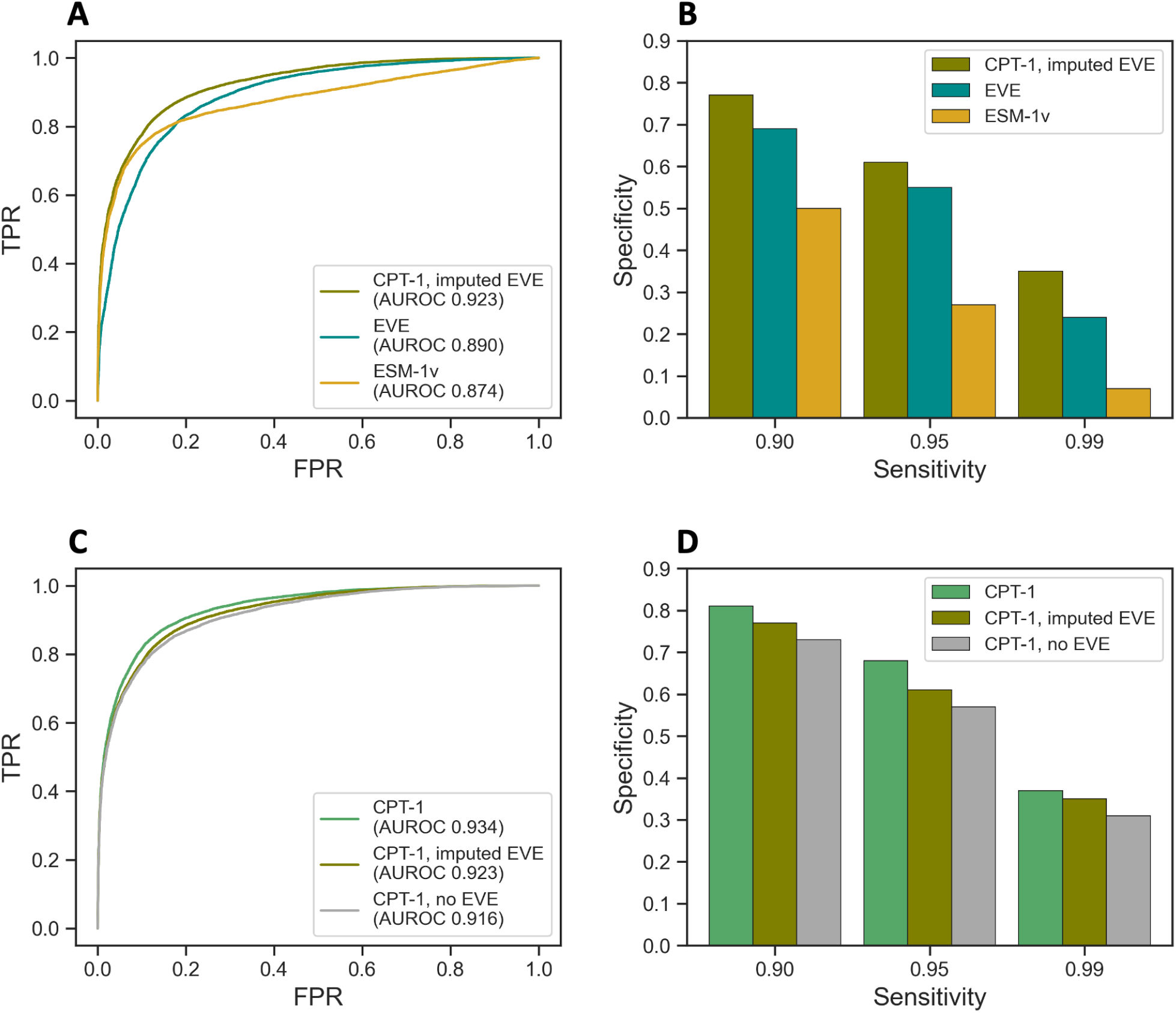
Cross-gene imputation. EVE scores are not available for the vast majority of human proteins. To scale our method to the whole human proteome, we imputed EVE scores and other features that depend on a large MSA in genes where they are not available. We assessed the quality of our imputation on genes where EVE scores are available, to measure how well we do compared to using the true values. (A-B) CPT-1 with imputed EVE still outperforms ESM-1v and the true EVE scores. (C-D) Imputed EVE scores improve performance of CPT-1 compared to removing them entirely, but there is still a gap to using the true EVE scores.

Using CPT-1 with imputed EVE, we were able to produce predictions for all missense variants in 90% of human genes. We used features based on the true EVE MSA in genes where this MSA was available and the imputed features in all other genes. In total, 3045 genes use the full CPT-1 model and 15,557 genes use CPT-1 with imputed EVE. The excluded 10% of genes were mostly due to not being contained in our vertebrate alignment dataset; many of these genes have not been clearly shown to produce a protein product.

## Discussion

The development of functional DMS assays and computational predictors have each been important to progress in missense variant effect prediction. We demonstrated that, although functional assays do not readily scale to the whole proteome directly, they can be a vital source of information for creating improved computational predictors. Using functional assay data of only five human proteins, we trained CPT-1, a computational predictor that significantly enhances the previous state-of-the-art. Our model is tested on a diverse set of proteins unseen during training and achieves improved performance by integrating vertebrate alignments and predicted structures with general protein sequence models. We used CPT-1 to release predictions for all missense variants in 90% of human genes.

We explored the integration of a larger set of functional assay datasets into our training scheme and found that this did not improve performance. A potential future direction is to develop a more powerful model architecture that may be able to better leverage this expanded data. Such a model could enable increased scope, such as modeling the effects of multiple mutants. Recent work has demonstrated progress modeling multiple mutants in the setting of functional assay prediction [33]. We found that vertebrate alignments provide strong signal for variant effect prediction that is non-redundant with EVE and ESM-1v. The utility of integrating vertebrate alignments across the human proteome points to exciting future directions. There are ongoing efforts to sequence a large number of vertebrate genomes [43]; as these data become available, more powerful models could be applied to deeper vertebrate alignments. Features calculated from AlphaFold2 structures also improve performance of our model. This result is interesting in light of the fact that AlphaFold2 primarily relies on the same evolutionary signals as EVE and ESM-1v to make structure predictions [30]. Recent work has discovered that AlphaFold2 has also learned an accurate representation of protein biophysics [44]. This additional signal may be responsible for the non-redundant information in AlphaFold2 structures.

CPT-1 mostly relies on general protein sequence variation models and sequence variation within vertebrates to predict the pathogenicity of missense variants. However, aspects of protein function that have emerged since the evolutionary divergence of vertebrates are still unlikely to be modeled well by CPT-1. Sequence variation may be insufficient to model such effects due to the sparsity of such data at very recent evolutionary timescales. Integrating experimental knowledge of the human protein interactome may help develop even more human-specific models [45], further increasing our understanding of various human diseases.

## Methods

### Datasets

We trained our models on data from the same functional assay on five human proteins, generated by the same research group [7,32]. These proteins are CALM1, MTHR, SUMO1, UBC9, and TPK1. The assay measures relative yeast fitness with different variants of the human protein of interest. We initially restricted ourselves to these proteins to ensure a high-quality, controlled training set while having enough diversity to transfer to entirely different proteins. The functional assay is also well-aligned with transferring to human clinical effects, since it measures the overall fitness of yeast as opposed to a specific biophysical property of the proteins. We additionally explored using functional assay data from 13 DMS experiments on human proteins from ProteinGym. These proteins are KCNH2, SCN5A, SC6A4, RASH, SYUA, PTEN, VKOR1, A4, P53, MSH2, TPOR, BRCA1, YAP1. These 13 were obtained by taking all human experiments from ProteinGym, removing all where EVE scores are not available, and only keeping the most recent experiment for each gene. We additionally excluded the dataset for gene TADBP because the distribution of variant effects was clearly not bimodal and ESM-1v has no significant predictive signal on this dataset. We found that adding these datasets to training did not increase performance of CPT-1 compared to our initial five high-quality and more homogeneous datasets (Supplementary Figure S2). We used this combined set of 18 experiments to assess performance on functional assay prediction. For the five main datasets, we trained functional assay prediction models on the other four proteins. For the additional 13 datasets from ProteinGym, we trained functional assay prediction models on all five main datasets.

We assessed our model for disease variant prediction on missense variants in ClinVar. We restricted to submissions with at least one star that were added since 2017, to ensure the dataset was high-quality. We additionally restricted to genes where EVE scores are available [8]. This left us with 24,155 variants in 1298 genes. We included variants annotated as “Benign” or “Likely Benign” and variants labeled as “Pathogenic” or “Likely Pathogenic” for our benign and pathogenic labels. We additionally compared our model to REVEL on genes that were not seen by REVEL at train time. For this comparison, we took our full dataset and removed any gene that had a one-star missense variant in ClinVar in 2017, with an annotation of “Benign”, “Likely Benign”, “Pathogenic”, or “Likely Pathogenic”. This left us with 3754 variants in 407 genes. Finally, in Supplementary Figure S1, we additionally restrict to the 50% of these variants with lowest allele frequency in gnomAD v2 [2]. This corresponds to a cutoff of 3.7 *×* 10^*−*4^.

We derived features from MSAs of orthologous sequences from 100 vertebrate and 30 mammalian species. These MSAs are available from the UCSC genome browser for most of the human genome, and were constructed using whole-genome alignment [28,29,34]. For some genes (228 in the EVE dataset), different isoforms were used in the vertebrate MSAs compared with other features, which are mainly based on UniProt. To resolve the discrepancy, we ran pairwise alignments between the vtMSA protein sequences and the UniProt sequences and only retained fragments of vtMSAs that can be aligned to UniProt sequences. We used the Bio.Align.PairwiseAligner implemented in the Biopython package with the setting: `model = ‘local’, match score = 5, mismatch score = -4, open gap score = -4, extend gap score = -0.5`.

We obtained predicted structures for all proteins from the AlphaFold2 human proteome database [31]. For proteins with a known experimental structure, the AlphaFold2 structure is generally highly accurate because the known structure has been provided as a template to AlphaFold2. Using all AlphaFold2 structures makes the input structures have a homogeneous format. We used contact statistics from AlphaFold2 structures for certain features and analyses. We extracted contacts using the Probe software [46], which notably identifies only sidechain-sidechain contacts.

### Features used in our transfer model

We initially considered a large set of potential features to include in our transfer model. Notably, we excluded predictors which were previously trained on clinical or functional assay data, to prevent data leakage. We also did not use features that do not extrapolate in an obvious manner to all 19 possible amino acid variants at a protein position (as opposed to all 9 possible single nucleotide variants in a codon). Our training functional assay data have a large amount of amino acid variants that are not expressible as single nucleotide variants. Restricting to full amino acid variant coverage allowed us to use more training data per protein and preserves applicability to functional assay prediction.

First, we included scores from the general protein homology models EVE and ESM-1v. We used the EVE scores as log-probabilities, rather than the final version which were normalized to a zero to one scale [8]. We obtained most EVE scores from the EVE dataset. We additionally computed EVE scores for SUMO1 and UBC9, which are part of our five training proteins but not included in the EVE dataset. By default, EVE does not provide scores at positions where the quality of the MSA is low; we imputed EVE scores at these positions using a within-gene *K*-nearest neighbors approach, which will be described in detail in the next section (see **Weighted KNN Imputation**). Imputed EVE predictions within a gene are less accurate than true EVE predictions but still increase the performance of our model (Supplementary Figure S3). We also generated predictions for human proteins outside of the EVE dataset where no EVE predictions were available. For these genes, we imputed EVE scores (and other features relying on the EVE MSA) using a cross-gene *K*-nearest neighbors approach (see **Weighted KNN Imputation**). Imputed EVE predictions across genes also increased the performance of our model (Figure 6).

We computed ESM-1v scores for all proteins in the UniProt collection of canonical transcripts for the human proteome (downloaded May 2022) [47]. We used the ESM-1v log probability difference to the wild-type amino acid, as in Meier *et al*. [9]. By default, ESM-1v does not accept proteins longer than 1022 amino acids. We developed a scheme to use ESM-1v on longer proteins using multiple sliding windows and used this scheme to compute ESM-1v scores for long proteins. Concretely, we calculated ESM-1v predictions with overlapping 1000 amino acid windows, with starting positions 250 amino acids apart on the protein sequence. Then for each mutation in the sequence, we used the score from the window whose center is the closest to the position of the mutation, as the center positions are expected to have better ESM-1v predictions given richer contextual information available.

To capture conservation at closer evolutionary timescales, we included features calculated using the 30-mammal and 100-vertebrate MSAs. Specifically, we obtained three types of frequencies for each mutation: the frequency of the wild-type amino acid at its position, the frequency of this mutant amino acid at this position, and the frequency of gaps in the alignment at this position. We refer to them as wild-type frequency, mutant frequency and gap frequency, respectively. These frequencies were log-transformed with offset 1. In genes where vtMSAs had different isoforms, certain regions were unmatched in pairwise alignment (see **Datasets**). Features in these regions were imputed with the weighted *K*-nearest neighbor (KNN) imputation strategy (see **Weighted KNN imputation**).

We included several features that explicitly use the AlphaFold2 structure of the protein. First, we calculated variant log-probabilities for all human proteins from three versions of ProteinMPNN, which was created with a focus on protein design [42]. Vanilla ProteinMPNN takes in protein structure with full protein backbone along with partial protein sequence. C*α* ProteinMPNN takes in protein structure with only alpha carbons for each residue along with partial protein sequence. C*α*-only ProteinMPNN uses only alpha carbons (no masked protein sequence). We normalized these scores as the log-probability difference to the wild-type allele log-probability, matching ESM-1v. Only Vanilla ProteinMPNN is used in CPT-1 after feature selection.

We also included two hand-designed structural features which we found perform well on functional assay data. These features aim to capture sequence variation in the EVE MSA conditioned on the structural environment around a position matching the environment in human proteins. Using the AlphaFold2 structure and the EVE MSA, we calculated for each position all other positions that form a sidechain-sidechain contact to it. Sidechain-sidechain contacts were calculated using the Probe software (see **Datasets**) [46]. Next, for each position, we filter the EVE MSA to sequences where the contact residues for that position have the same amino acids as in the human sequence. However, we only use the two contact residues where the human amino acid appears most frequently in the EVE MSA, to keep the number of sequences from becoming too small. We additionally only allowed conditioning on residues with pLDDT greater than 70 in the AlphaFold2 structure, and residues with pLDDT less than 70 did not have any conditioning used. Finally, for each position, we compute the frequency of the human allele and all possible alternative amino acids in the filtered MSA. These features are the *conditioned wild-type score* and *conditioned mutant score*, respectively.

Some human proteins have multiple fragment AlphaFold2 structures in the AlphaFold2 human proteomes. For these proteins, we computed structural features using the fragment that maximized the pLDDT of that position. We also included sidechain-sidechain contact count and AlphaFold2 pLDDT as features.

Finally, we included amino acid descriptors, which are featurizations of amino acids that encode properties such as charge, polarity, hydrophobicity, size, and local flexibility [48]. The descriptors we used include Cruciani properties [49], VHSE [50], Z-scales [51], ST-scales [52], ProtFP [53], and Georgiev’s BLOSUM indices [54]. We used the differences in the descriptor values between the mutant amino acid and the wild-type amino acid as features for each mutation.

### Weighted KNN imputation

We used a strategy based on *K*-nearest neighbor imputation to impute missing values in the feature matrix. Take the EVE scores as an example. To train a KNN model on a given gene, we first calculated the Spearman correlation between each feature and the EVE scores at the available mutations within the gene. Then the five most highly correlated features together with the EVE scores were used to build the KNN model. When calculating the distance matrix, each feature was weighted by its correlation value with the EVE scores, which was implemented as scaling each of the features by the correlation value after standardization. EVE scores were assigned weight 1 in the scaling.

When applying the fitted model to impute missing values, we used two strategies in this study, which we refer to as within-gene imputation and cross-gene imputation. For genes included in the EVE dataset, we used within-gene imputation to directly impute missing EVE scores with the model fitted on that gene. However, for genes not included in the EVE dataset where no EVE score is available for any mutation, we first fit five KNN models on the five training genes, used them to impute the EVE scores for all the mutations and then averaged the outputs across the five models to get the imputed values.

The within-gene imputation strategy was also used to impute missing vtMSA features. The cross-gene imputation strategy was used to impute missing structure-conditioned scores. We used the implementation of the KNN model in the python sklearn package (sklearn.impute.KNNImputer) with the number of nearest neighbors as 10 leaving other parameters as default.

### Model architecture and training

Our models were set up as either logistic regression to classify “functionally normal” from “functionally abnormal” mutants (for clinical disease variant prediction) or as linear regression to predict functional assay score (for hold-out protein functional assay prediction). We trained a separate linear model for each training protein and ensembled them by averaging the model predictions at test time; we found this to be an effective method to adjust for batch effects across each protein. We found that more complex, non-linear models did not transfer to held-out proteins well. We analyzed the impact of using variable numbers of training proteins (Supplementary Figure S2). Benefits appear to be saturating at all five proteins used; additional proteins may be more useful if diversity is increased.

Although functional assay data for our five training proteins was generated by the same research group with almost the same protocol, the distribution of scores varies significantly between proteins (Supplementary Figure S6). To remedy this, and because pathogenicity annotations are binary, we decided to binarize the functional assay scores to train the classification model for disease variant prediction. Specifically, we standardized the data by taking the top 40% of variants from each protein as functionally normal and the bottom 40% as functionally abnormal. We found that this percentile-based binarization provided stable results, and these results did not depend much on the exact binarization threshold (Supplementary Table S4).

We used a global feature rescaling for all proteins, calculated from our five training proteins. We scaled the features to unit standard deviation, but calculated these standard deviations by reweighting all samples from each training protein to total weight one, so that each protein has the same total weight in calculating rescaling weights. This prevents the rescaling terms from being biased towards larger proteins in our training dataset. The MSA features are frequencies and do not behave well under the scaling and were therefore left as is.

We initially considered a large list of candidate features and employed feature selection to reduce this list before fitting linear models. We used average AUROC from cross-validation on the training functional assay data as performance metrics to select features (for regression, we instead use Spearman correlation). Specifically, in each fold we leave out one protein in the training set as the validation set. We use the remaining proteins to fit the model and then evaluate the performance using the validation protein. For the disease variant classification model, a 5-fold cross-validation was performed for all 5 training proteins. For the functional assay score regression model, a 4-fold cross-validation was performed excluding the held-out protein. We always include the two protein homology models as features, i.e. ESM-1v and EVE scores. For the remaining features, we used a two-step scheme for feature selection: the first step selects features from the 100-vertebrate MSA, 30-mammal MSA, ProteinMPNN and structure-conditioned score categories and the second step selects amino acid descriptor features. In the first step, we exhaustively searched all combinations of features within each of the four categories above, restricting that at least one feature was selected from each category. Then in the second step, given the large number of amino acid descriptor features, we used forward selection to reduce the computational burden. To go through the procedures, we start with the two features of ESM-1v and EVE. Then in the first step, we add the set of 100-vertebrate MSA features that achieves the best performance on the validation protein. Then we add the best sets of 30-mammal MSA features, ProteinMPNN features and structure-conditioned score features in the same way. Then in the second step, we greedily select the first few amino acid descriptor features that make the largest improvements in performance. Final selected features for CPT-1 are reported in Supplementary Table S1. To examine the effects of removing vertebrate alignments, 100-vertebrate and 30-mammal mutant frequencies were removed and models were retrained. To examine the effects of removing structure, ProteinMPNN and conditioned mutant/wild-type frequencies were removed and models were retrained.

## Data and Code Availability

Codes and data for training the CPT-1 model and reproducing main results are available at https://github.com/songlab-cal/CPT. We release pre-computed CPT-1 predictions for all missense variants in 90% of human genes at https://doi.org/10.5281/zenodo.7954657. We also created an interactive app for visualizing and downloading CPT-1 predictions for individual proteins https://huggingface.co/spaces/songlab/CPT.

## Competing Interest Statement

The authors declare no competing interests.

## Acknowledgements

We would like to thank Sanjit Batra, Gonzalo Benegas, Chloe Hsu, Sergey Ovchinnikov, Junhao Xiong, and members of the Song Lab for helpful discussion. This research is supported in part by an NIH grant R35-GM134922, a grant from the Koret-UC Berkeley-Tel Aviv University Initiative in Computational Biology and Bioinformatics, and a grant from the Noyce Initiative UC Partnerships in Computational Transformation Program.

## Notes

### Competing Interest Statement

The authors have declared no competing interest.

### Summary of Updates

We have carried out extensive further analyses to test the performance of our model. In particular, we have explored the possibility of training on a larger collection (eighteen) of DMS datasets. Also, we have added a benchmarking study against REVEL.

