## Supplementary Tables ad Figures for "Cross-protein transfer learning substantially improves disease variant prediction"

This file contains

- Supplementary Figures S1-S6
- Supplementary Tables S1-S4

[*]These authors contributed equally to this work.

#### Supplementary Figures

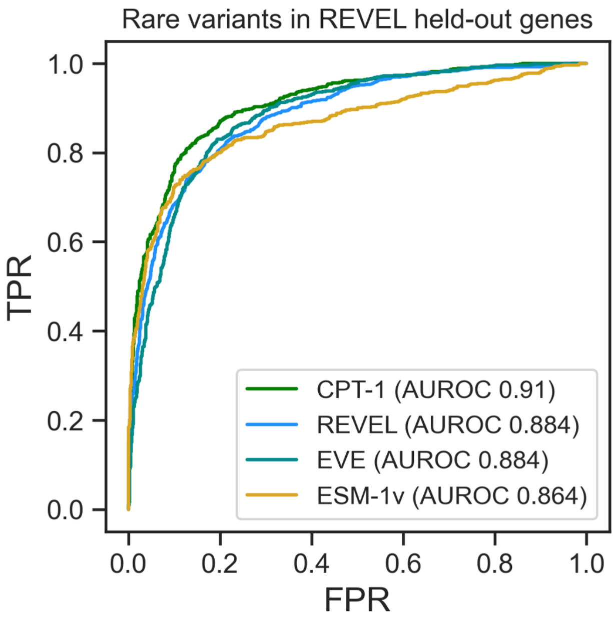


Figure S1: **Comparison with REVEL on rare variants.** CPT-1 outperforms REVEL on genes not seen by REVEL at train time (Figure 2). If we additionally restrict to the 50% of variants with lowest allele frequency (a cutoff of $3.7 \times 10^{-4}$ in gnomAD), the margin over REVEL becomes larger. Also, REVEL does not outperform EVE in this rare variant subset.

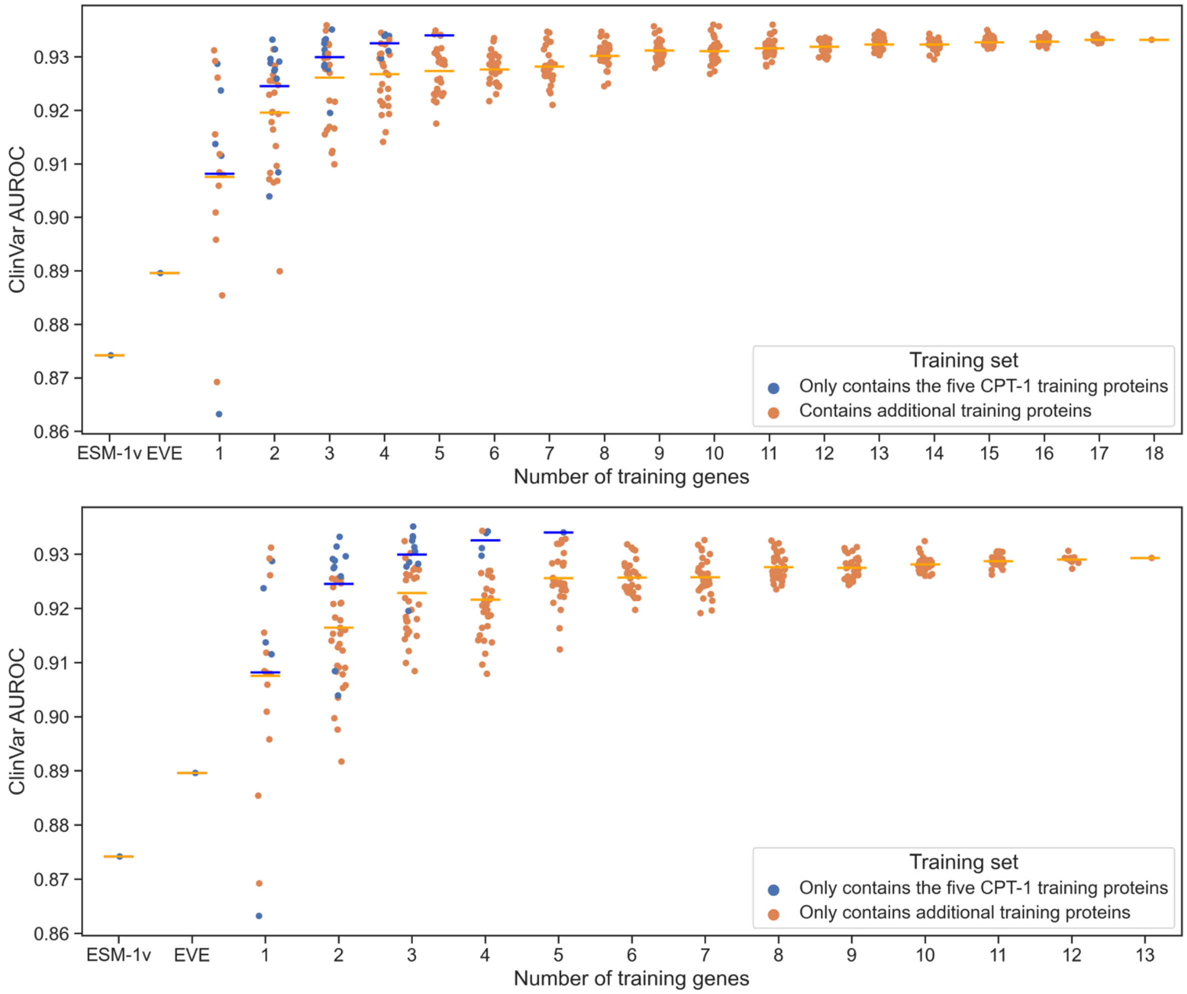


Figure S2: **Impact of choice of training genes.** CPT-1 is trained on functional assay data from five proteins generated by the same lab. We also examined the possibility of training on additional datasets from ProteinGym. Each dot indicates a specific choice of training genes, with the mean shown as a horizontal bar. (Top) Selecting from only the five main datasets yields better performance compared to a random, equal number from the full collection of 18 datasets. In addition, using all datasets does not outperform only using the five main datasets. (Bottom) Selecting from only the five main datasets yields better performance compared to a random, equal number from the 13 proteins from ProteinGym. These results demonstrate the value of using DMS data from the same assay and lab.

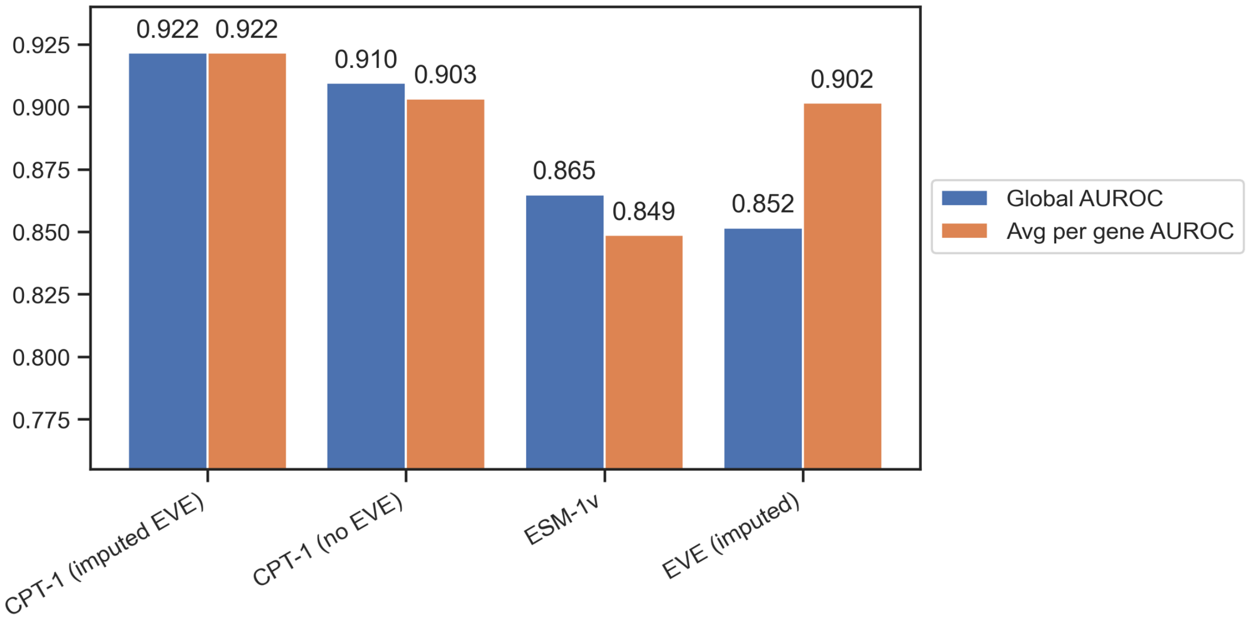


Figure S3: **AUROC results for positions where within-gene imputation of EVE predictions was performed.** EVE declines to make variant effect predictions at positions with low alignment quality in the EVE MSA. We imputed EVE predictions at these positions using a nearest-neighbors approach within the gene. Imputed predictions increase CPT-1 performance both in terms of overall AUROC and per-gene AUROC. ESM-1v performs worse on these low-alignment quality variants as well, compared to ESM-1v performance on other variants.

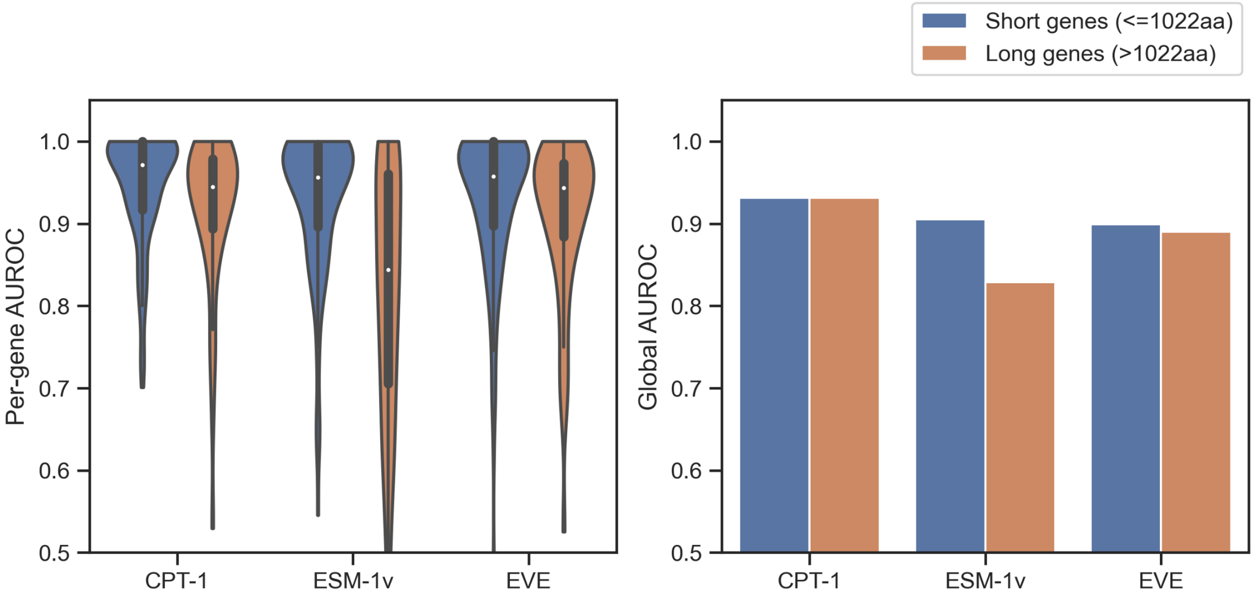


Figure S4: **Extending ESM-1v to long genes**. ESM-1v by default does not accept proteins longer than 1022 amino acids. To enable full coverage of human proteins, we developed a scheme to apply ESM-1v to longer proteins. ESM-1v with this extension performs worse on long genes, but CPT-1 does not suffer a performance drop. EVE performs as well on long proteins as on short.

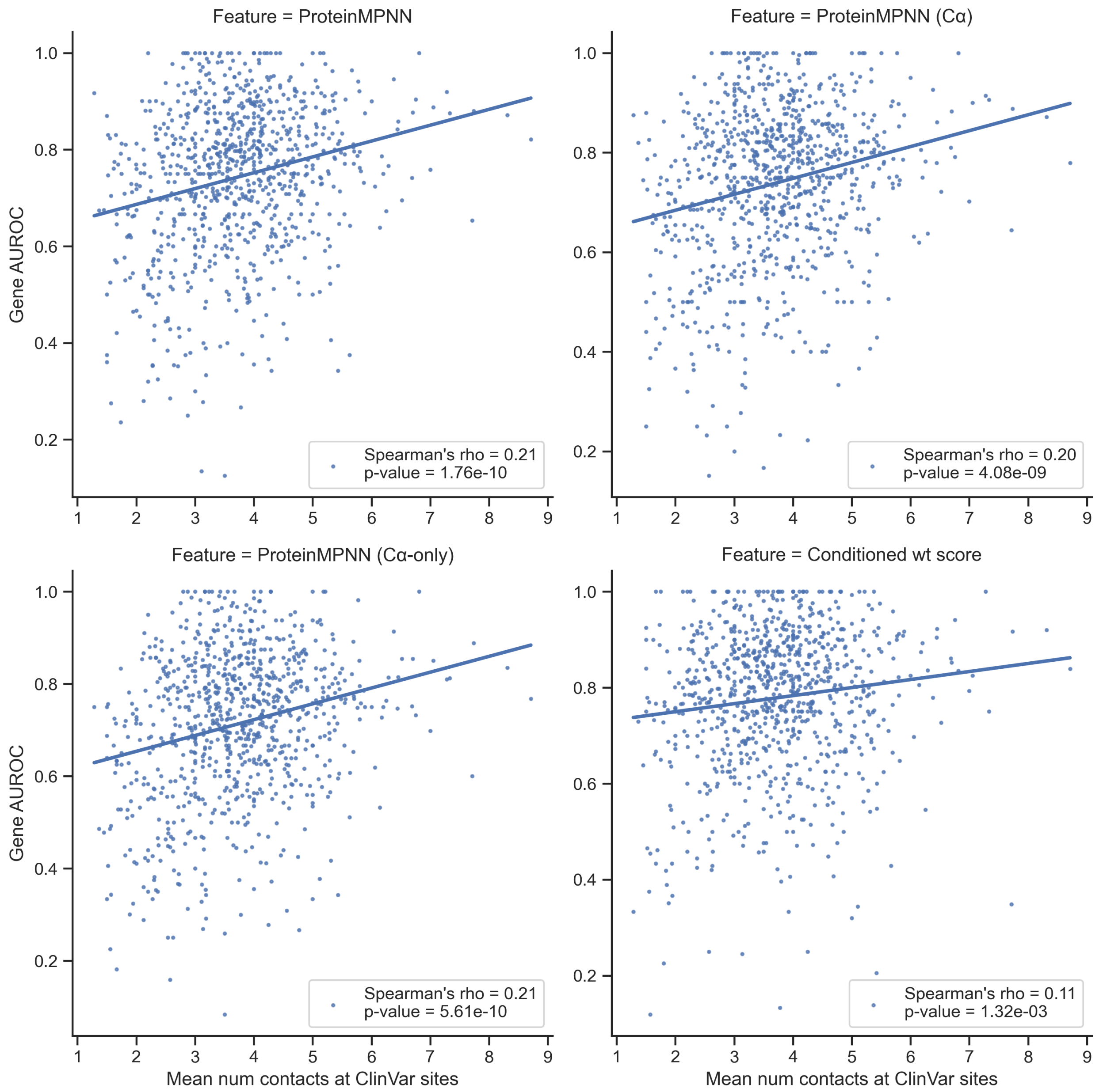


Figure S5: **Contact counts**. We calculated the number of sidechain-sidechain contacts formed by the position of each ClinVar variant and took the average of these counts in each protein. Structural features perform better in genes where ClinVar variant positions have more contacts. Contact count is itself a modest predictor of variant pathogenicity (0.69 AUROC), indicating that ClinVar variants in the core of proteins are more likely to be pathogenic.

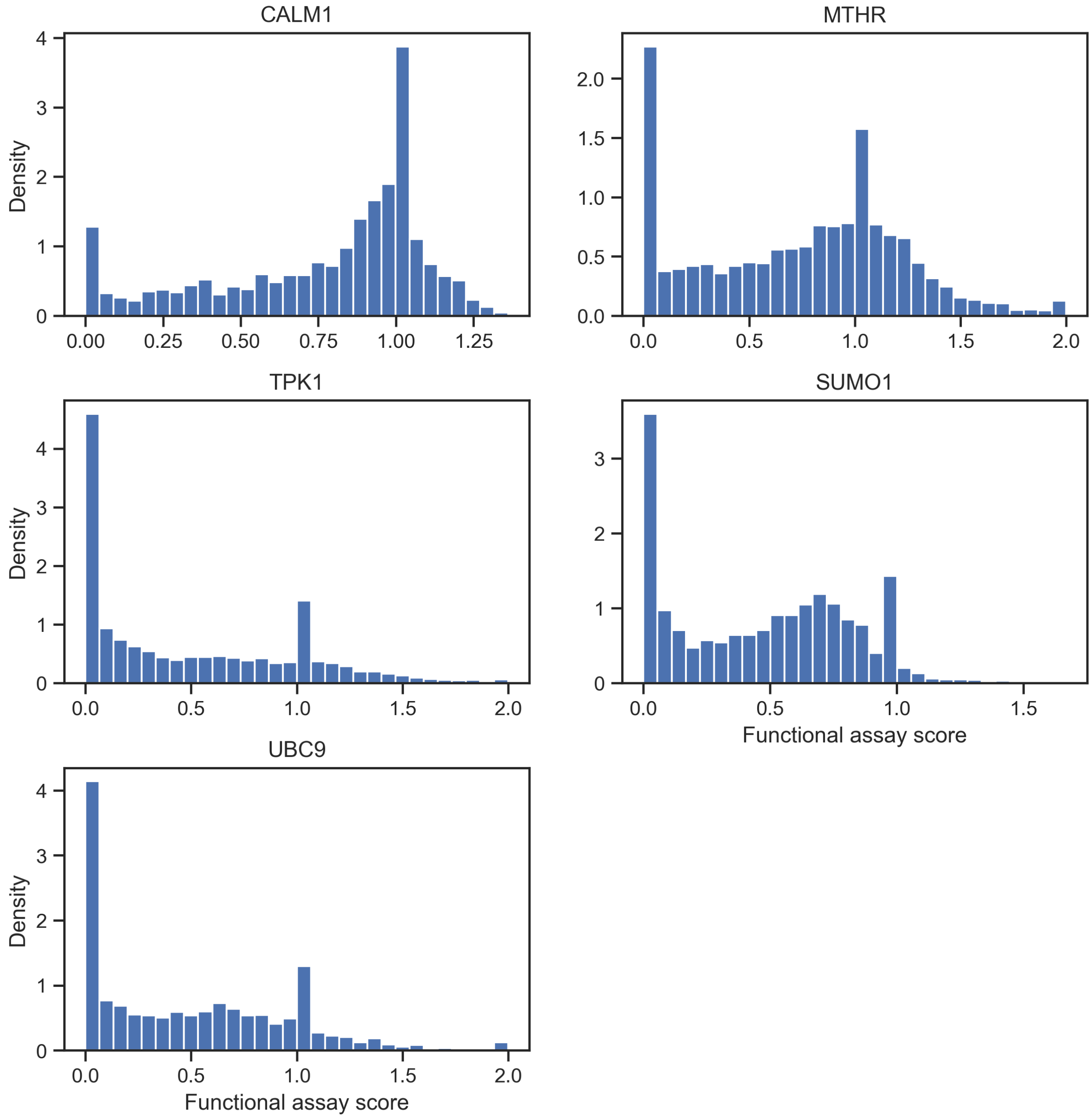


Figure S6: **Distribution of functional assay scores**. Functional assay scores for our five training proteins were generated by the same group with a standardized scale. A score of 0 is set to match the average effect of a nonsense (stop-gain) variant, while a score of 1 is set to match the average effect of a synonymous variant. Some datasets were truncated at 0, so we truncated all 5 at 0. Despite the relative homogeneity between these datasets, the distribution of variant scores differs significantly between genes, which led us to use binarization for our disease variant model.

### Supplementary Tables

| Feature | Type | Description | Source |
|---|---|---|---|
| EVE evolutionary index | General sequence variation | Scores from neural model of protein sequences in MSA | Frazer *et al.*, 2021 [1] |
| ESM-1v | General sequence variation | Scores from language model of protein sequences | Meier *et al.*, 2021 [2] |
| 100 vertebrate mut. freq. | Vertebrate alignment | Mutant frequency in orthologous proteins from 100 vertebrates | UCSC Genome Browser [3, 4] |
| 30 mammal mut. freq. | Vertebrate alignment | Mutant frequency in orthologous proteins from 30 mammals | UCSC Genome Browser [3, 5] |
| Conditioned wildtype score | Structure + Sequence variation | Wildtype freq. in EVE MSA, conditioned on contact residues matching human protein | Novel |
| Vanilla ProteinMPNN | Structure | Scores from neural network on protein structure and partial sequence | Dauparas *et al.*, 2022 [6] |
| BLOSUM Index 1 | Amino Acid Descriptor | The first component of BLOSUM62-derived indices. | Georgiev, 2009 [7] |
| ST6 | Amino Acid Descriptor | The sixth principal component of ST-scales. | Yang *et al.*, 2010 [8] |
| VHSE6 | Amino Acid Descriptor | The sixth principal component (electrical properties) of VHSE-scales. | Mei *et al.*, 2005 [9] |

Table S1: **Final selected features for CPT-1**. CPT-1 is trained on features based on general sequence variation, sequence variation in vertebrates, protein structure, and amino acid descriptors. Certain features are imputed across genes for the whole-proteome scale version of CPT-1 (underlined). For all amino acid descriptor features, the difference from wild-type to mutant amino acid is used.

| Model | Overall AUROC | Avg gene AUROC | Median gene AUROC | Spec @ 95% Sens | Sens @ 95% Spec |
|---|---|---|---|---|---|
| CPT-1 | 0.934 | 0.933 | 0.958 | 0.675 | 0.698 |
| CPT-1 (no vtMSA) | 0.925 | 0.925 | 0.951 | 0.607 | 0.693 |
| CPT-1 (no structure) | 0.93 | 0.932 | 0.958 | 0.650 | 0.696 |
| CPT-1 with imputed EVE | 0.923 | 0.918 | 0.940 | 0.611 | 0.662 |
| EVE | 0.89 | 0.919 | 0.949 | 0.548 | 0.512 |
| ESM-1v | 0.874 | 0.877 | 0.931 | 0.273 | 0.640 |
| 100vt wt frequency | 0.865 | 0.865 | 0.890 | 0.508 | 0.00 |
| Vanilla ProteinMPNN | 0.781 | 0.765 | 0.791 | 0.134 | 0.390 |
| Conditioned mt score | 0.826 | 0.844 | 0.881 | 0.287 | 0.435 |
| Conditioned wt score | 0.719 | 0.807 | 0.819 | 0.196 | 0.185 |

Table S2: **Detailed classification performance on ClinVar missense variants.** CPT-1 outperforms baselines on all metrics. Per-gene ClinVar AUROC is calculated in 274 genes with at least four benign and four pathogenic ClinVar variants. The margin over EVE and ESM-1v depends heavily on vertebrate alignments and, to a lesser extent, on structure. The cross-gene imputation version of CPT-1 is used to produce predictions at whole-proteome scale; it performs worse than using true feature values, but outperforms EVE and ESM-1v. 100-vertebrate MSA wild-type frequency is a powerful baseline in the high-sensitivity regime. Structural features perform worse alone but have enough non-redundant signal to be worth including.

| Protein | CPT-1 | EVE | ESM-1v |
|---|---|---|---|
| Average | **0.448** | 0.380 | 0.412 |
| SUMO1 | **0.640** | 0.606 | 0.580 |
| UBC9 | **0.584** | 0.574 | 0.558 |
| TPK1 | 0.374 | 0.314 | **0.396** |
| MTHR | **0.330** | 0.293 | 0.314 |
| CALM1 | 0.301 | 0.291 | **0.318** |
| SC6A4 | **0.567** | 0.520 | 0.542 |
| YAP1 | 0.560 | **0.617** | 0.441 |
| P53 | **0.555** | 0.452 | 0.553 |
| BRCA1 | **0.555** | 0.515 | 0.450 |
| VKOR1 | **0.534** | 0.464 | 0.474 |
| PTEN | **0.507** | 0.360 | 0.454 |
| RASH | 0.467 | **0.480** | 0.405 |
| A4 | **0.456** | 0.343 | 0.435 |
| TPOR | **0.439** | 0.352 | 0.400 |
| MSH2 | **0.417** | 0.394 | 0.400 |
| KCNH2 | **0.358** | 0.023 | 0.295 |
| SYUA | **0.304** | 0.156 | 0.267 |
| SCN5A | 0.120 | 0.082 | **0.135** |

Table S3: **Detailed regression performance on DMS datasets of human proteins.** Spearman's $\rho$ of CPT-1, EVE, and ESM-1v are shown for eighteen DMS datasets of human proteins. The top five are the CPT-1 training proteins and the remaining thirteen are from ProteinGym. The highest performing method for each protein is bolded.

| Model | Overall AUROC | Avg gene AUROC |
|---|---|---|
| CPT-1 (30, 50 binarize) | 0.930 | 0.928 |
| CPT-1 (30, 60 binarize) | 0.931 | 0.928 |
| CPT-1 (30, 70 binarize) | 0.930 | 0.927 |
| CPT-1 (40, 50 binarize) | 0.933 | 0.929 |
| CPT-1 (40, 60 binarize) | 0.934 | 0.934 |
| CPT-1 (40, 70 binarize) | 0.933 | 0.930 |
| CPT-1 (50, 50 binarize) | 0.933 | 0.930 |
| CPT-1 (50, 60 binarize) | 0.934 | 0.933 |
| CPT-1 (50, 70 binarize) | 0.933 | 0.931 |
| ESM-1v | 0.874 | 0.876 |
| EVE | 0.890 | 0.919 |

Table S4: **CPT-1 performance is stable under different choices of training data binarization.** We binarized training functional assay data for CPT-1 based on percentile within each protein; variants with score below the 40th percentile were labeled pathogenic, and variants with score above the 60th percentile were labeled benign. Changing these thresholds does not change performance on ClinVar variants much, and CPT-1 always outperforms baselines.
